# Hair cell population integrity necessary to preserve vestibular function

**DOI:** 10.1101/2025.03.19.644154

**Authors:** Louise Schenberg, François Simon, Aïda Palou, Cassandre Djian, Michele Tagliabue, Jordi Llorens, Mathieu Beraneck

## Abstract

Vestibular dysfunction constitutes a major medical concern, and regeneration of hair cells (HC) is a primary target of gene therapy aimed at restoring vestibular functions. Thus far, therapeutic trials in animal models targeting vestibular loss associated with genetic diseases have yielded variable and partial results, and the functional identity and quantity of HCs required to restore minimal or normal vestibular function remain undefined. Indeed, direct comparisons between structural pathology and quantitative assessments of vestibular dysfunctions are lacking in humans and are rather limited in animal models, representing a significant gap in current knowledge. Here, we present an innovative methodology to bridge the gap between HC integrity and functional vestibular loss in individuals. Gradual vestibular deficits were induced through a dose-dependent ototoxic lesion, quantified with canal or utricular-specific vestibulo-ocular reflex tests, and were then correlated in all individuals with the loss of type I and type II HCs in different regions of ampulla and macula. Our findings reveal that the structure-function relationship is nonlinear, with lower bound of approximately 50% of HCs necessary to retain minimal vestibular function, and threshold exceeding 80% to preserve normal function, thus shedding light on population coding mechanisms for vestibular response. Our data further support the decisive role of type I, rather than type II, HC in the tested VOR functions.

## INTRODUCTION

The vestibular system is fundamental to the ability of vertebrates to detect head movements and to discriminate between self-motion and environmental motion. Angular and linear head accelerations are encoded by two distinct sets of inner ear organs: the three semi-circular canals, which encode rotational movements across all planes, and the otolithic organs, consisting of the saccule and utricule, which encode head accelerations relative to gravity (Angelaki and Cullen, 2008). Vestibular information is conveyed from the inner ear to the brainstem vestibular nuclei and vestibulo-cerebellum, where it is processed for various functional pathways. Thus, the vestibulo-ocular pathway is responsible for gaze stabilization, the vestibulo-spinal pathway for posture and balance, and the vestibulo- thalamic pathways for orientation and self-motion perception by providing information to subcortical and cortical circuits. Vestibular dysfunction constitutes a major medical concern, as one of its manifestations, dizziness, affects 15–35% of the general population, with a prevalence rate of up to 85% in individuals over the age of 80 (Agrawal et al., 2013). Age-related alterations of both vestibular function and the integrity of vestibular hair cells have been reported in humans (Merchant et al., 2000; Rauch et al., 2001; Lopez et al., 2005; Walther and Westhofen, 2008).

Despite its broad involvement in numerous vital functions, the initial steps of vestibular encoding remain incompletely understood and the topic of ongoing research. Vestibular sensory transduction depends on mechanoreceptors known as hair cells (HC) located in the neuroepithelium of the different vestibular organs. These cells are characterized by their apical filiform extensions, the stereocilia, projecting from the epithelium into the lumen of the labyrinthic cavities. The HCs convert the mechanical movements induced by head motion on the stereocilia bundle into electrochemical signals for sensory input into the vestibular nerve. Vestibular HCs in mammals are categorized into distinct subtypes, each with unique morphological, molecular, electrophysiological and synaptic characteristics (Lysakowski and Goldberg, 1997; Rüsch et al., 1998; Eatock and Songer, 2011; Eatock, 2018; McInturff et al., 2018). Type I HC found in amniotes (reptiles, birds and mammals) (Mackowetzky et al., 2021) are enveloped by a calyx-shaped terminal from a vestibular afferent neuron. These postsynaptic calyces have recently been shown to process exceptionally rapid (0.3ms) transmission through direct electrical coupling between the hair cell and its afferent (Eatock, 2018), alongside classical quantal/glutamatergic communication (Contini et al., 2024). The Type I/calyceal synapse appears particularly suited for the rapid information processing, probably essential to feed the short-delay reflexive vestibular pathways such as the vestibulo-ocular reflex (5-7ms, Huterer and Cullen, 2002). Therefore, the encoding of rapid head motions with high frequency content, as observed during natural head activity (Carriot et al., 2017), may primarily depend on the Type I HC transducing transient, rather than sustained, stimuli (Songer and Eatock, 2013). This information would be preferentially conveyed by one of the two types of afferents defined according to electrophysiological criteria: the irregular afferents that preferentially reach the apex/center of the crista ampullaris and the central striola region of the maculae of the otolith organs (Curthoys et al., 2017).

Type II HC exhibit the greatest similarity to those observed in the cochlea and in non-mammalian vertebrates and are believed to be evolutionarily more ancient (Maudoux et al., 2022). Characterized by thinner and fewer stereocilia per hair bundle, they can be identified by their cylindrical-shaped cell body, which connects to multiple afferent bouton synapses. Type II HC are also predominantly associated with electrophysiologically defined regular afferents, more prevalent in peripheral and non-striolar zones of the different vestibular organs (Goldberg, 2000). Type II HC transmit sensory information exclusively through classical neurotransmitter release to their afferents, and are often considered as particularly well suited to process low-frequency, tonic, and regular activity. Despite the absence of direct correspondence between type II hair cells and regular afferents, these likely constitute the two canonical presynaptic and postsynaptic elements of the peripheral pathway responsible for encoding sustained stimuli, with lower detection thresholds (Jamali et al., 2016).

Loss of inner ear or vestibular HCs frequently occurs as a secondary ototoxic effect of aminoglycoside antibiotics (streptomycin, neomycin and gentamicin), and chemotherapy agents such as cisplatin (Halmagyi et al., 1994; Selimoğlu et al., 2003; Wu et al., 2021). In addition, studies have reported age-related changes in vestibular function (Paplou et al., 2021) as well as structural alterations in vestibular HCs in humans (Rauch et al., 2001; Lopez et al., 2005; Walther and Westhofen, 2008; Merchant et al., 2000). However, direct correlations between structural pathology and quantitative assessments of vestibular dysfunction remain rare in humans and are relatively limited in animal models (Maroto et al., 2021; Paplou et al., 2021). Despite significant development in understanding the role of the different HCs in vestibular processing, the precise relationship between HC and the resultant functional deficits remains poorly defined (Fleihan et al., 2024).

The inner ears of many non-mammalian species (e.g amphibians, reptiles, birds) are able to produce HCs throughout their lifespan (Elliott et al, 2018). In contrast, in the mature mammalian ear, only a fraction of vestibular type II, and not type I, HCs can regenerate into an immature form from differentiating supporting cells in the extrastriolar/peripheral regions (Golub et al., 2012; Sayyid et al., 2019; Wang et al., 2019). Several animal studies have attempted to investigate the restoration of vestibular function through gene therapy or pharmacological interventions aimed at promoting HC regeneration (Emptoz et al., 2017; Lahlou et al., 2024). However, these trials have produced inconsistent and often partial outcomes, largely due to the complexity of vestibular signal processing and the heterogeneous nature of HC loss. Overall, HC regeneration remains a central goal in efforts to restore vestibular and cochlear functions (Emptoz et al., 2017).

Fundamental questions regarding the functional significance of vestibular HC persist. First, the requisite quantity or relative proportion of vestibular HC restoration necessary to achieve specified levels of functional recovery remains unknown. Second, the functional importance of the vestibular HC (e.g. type I and type II) located within various vestibular organs and epithelial regions (central/striolar zone versus periphery/extra-striolar zone) requires better characterization.

To address these gaps, we employed an innovative approach to establish a precise correlation between HC integrity and vestibular function in individual rodents (Maroto et al., 2021). In our study, vestibular deficits were gradually induced using a variety of doses of an ototoxic compound. The extent of vestibular dysfunction was quantified through canal- and utricular-specific vestibulo-ocular reflex (VOR) tests, allowing for a robust and quantitative assessment of vestibular performance. By systematically correlating these functional measurements with histological analyses of type I and type II HC loss in distinct regions of the ampullae and maculae, we aimed to elucidate the structure-function relationship governing the vestibulo-ocular reflexes.

Our findings reveal that vestibular function exhibits threshold-dependent characteristics, with a lower bound of approximately 50% hair cell preservation required to sustain minimal vestibular function. In contrast, the preservation of normal vestibular function is reached at a threshold of around 80%. These insights contribute to our understanding of population-coding strategies within the vestibular system and provide a framework for future research investigating type-specific HC regeneration. Our data further support the pivotal role of type I HC in mediating VOR functions, as their depletion was more strongly associated with vestibular deficits compared to the loss of type II HC (Schenberg et al., 2023).

## RESULTS

### Dose-dependent effects of the treatment on the canal- and otolith-dependent VOR

To investigate the dose-response effect of the ototoxic substance 3,31-iminodiproprionitrile (IDPN), we performed in adult C57Bl6/J mice a single IP injection of IDPN at increasing doses: 0 (Control, n=7), 16 (n=8), 24 (n=8), 32 (n=7) and 40 (n=6) mmol/kg. Organ-specific vestibular function was quantified using gaze stabilizing reflexes tests before the injection, then 2 and 3 weeks after. There was no significant difference between vestibular function observed at D14 and D21 (mixed model analysis, see Supplementary Figure 1), confirming that the IDPN-induced toxicity is stable after 2 weeks(Llorens et al., 1993; Soler-Martin et al., 2006). To reduce measurement noise, data from D14 and D21 are pooled in the rest of the manuscript.

Horizontal canal function was assessed with two complementary tests. First, horizontal sinusoidal rotations at different frequencies were performed in the dark to record the angular horizontal vestibulo-ocular reflex (aVOR) the gain of which are shown in the bode plots of Figure 1.A for the different groups and tested frequencies. IDPN exposure at different doses (abbreviated as [X]mmol/kg) leads to a significant decrease of the aVOR gain for concentrations above [16] (Interaction Dose x Frequencies F(16, 124)=15.4, p<0.0001, see statistics table 1). The aVOR gain of the [24] group is decreased by ∼50% compared to the Control group ([Control] vs [24]; at 0.2Hz, p=0.0625; 0.5Hz, p=0.0147; 0.8Hz, p=0.0351; 1Hz p= 0.0454 and 2Hz p=0.0261) and is significantly reduced by >90% at all frequencies for the 2 highest doses of [32] and [40] (p<0.001). The second test, Hsteps, consisting in a single abrupt horizontal acceleration (peak acceleration 500°/s²; Figure 1.B) shows a significant decrease of gain similar to aVOR for the 3 highest doses (F(4, 31)=28.58, p<0.0001; [Control] vs [16] p=0.9956; [Control] vs [24], [Control] vs [32] and [Control] vs [40], p<0.001). While the [32] and [40] groups have no residual response, the [24] group show an intermediate VOR loss ([16] vs [24] p<0.001; [24] vs [32] p=0.0485).

**Figure 1.**
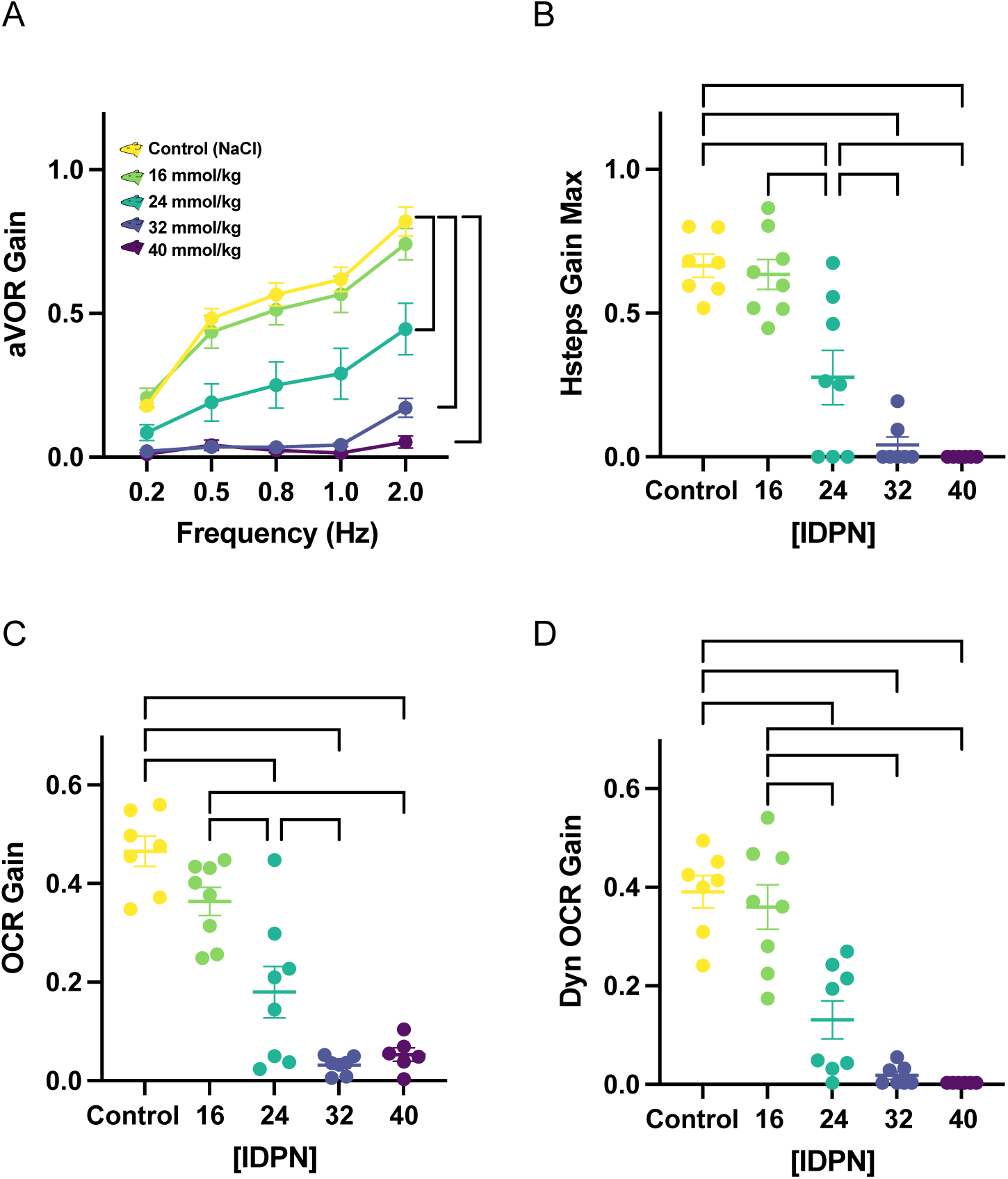
Dose-dependent ototoxic effect on vestibulo-ocular reflexes. A) Bode plot of the gain of the horizontal angular vestibulo-ocular reflex at different frequencies (tested with sinusoidal rotations at peak velocity 30°/s) for the 5 different concentration tested. B) Maximal gain response in respond to abrupt horizontal acceleration/deceleration (steps of acceleration from 0 to 50°/s, with peak velocity 250°/s²). C) Gain of the ocular-counter roll reflex measured during static roll in range ±0-40. D) Dynamic ocular-counter roll reflex gain. The gain was calculated using the vertical component of the compensatory eye movement evoked in response to off-vertical axis rotation with a tilt angle of 17°. In all panels, sample sizes are: [Control] (n=7), [16] (n=8), [24] (n=8), [32] (n=7) and [40] (n=6). Vertical and horizontal brackets indicate significant differences (p<0.05). Values represented mean±SEM.

**Table 1:**
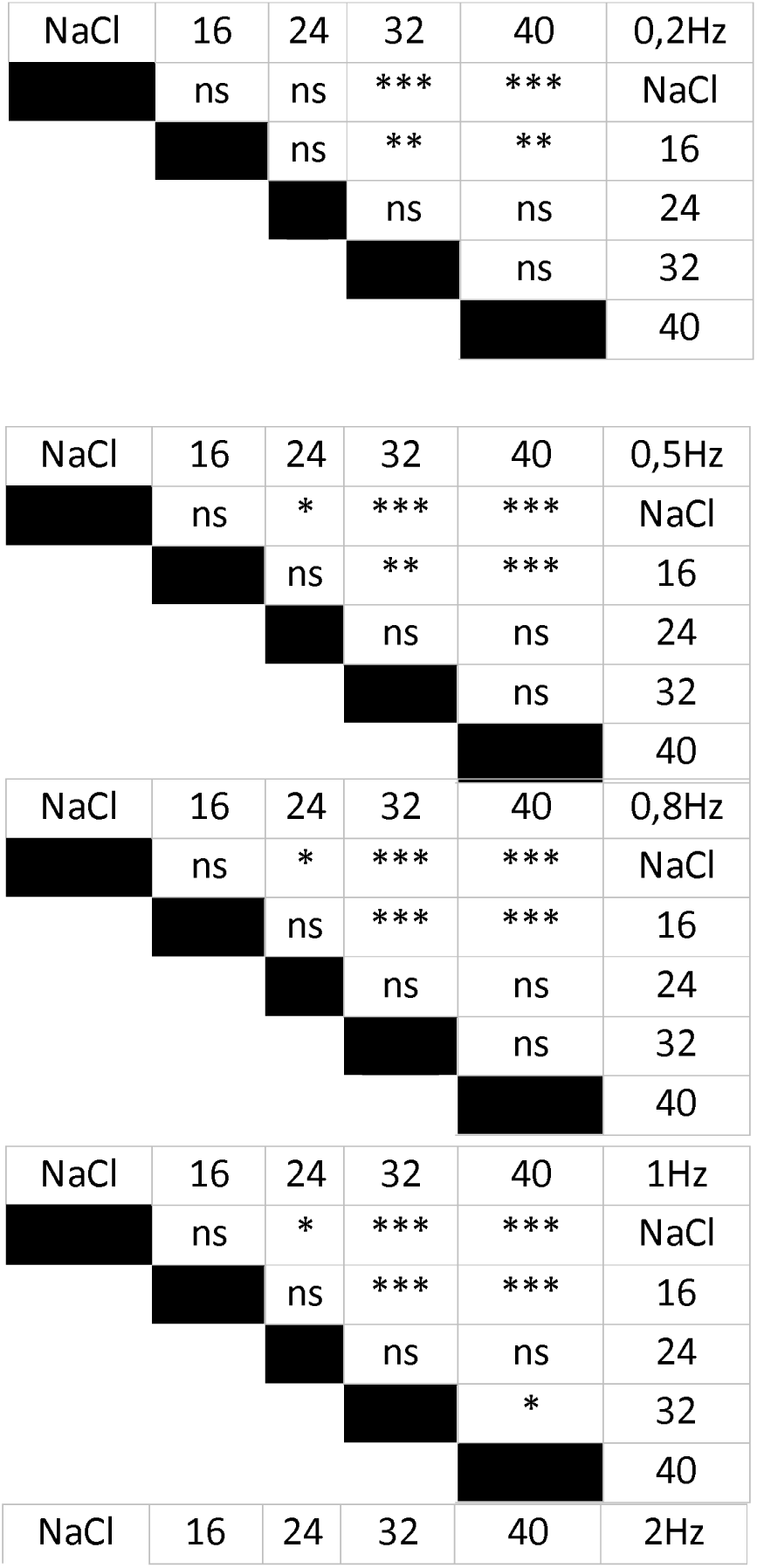
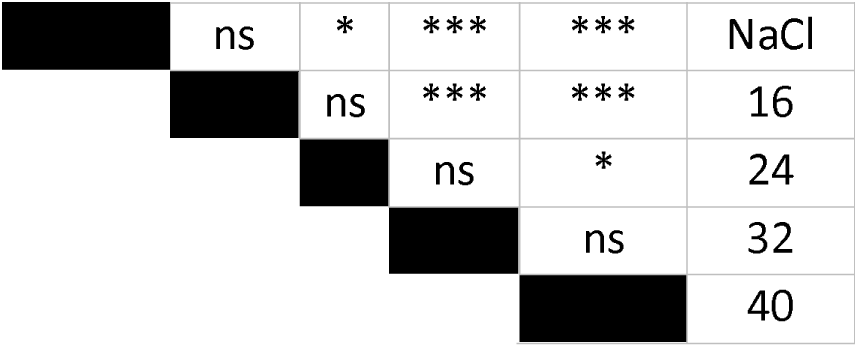
Statistical tables for the 5 frequencies tested of the aVOR (Figure 1)

Specific otolithic (utricular) function was quantified using both static ocular counter-roll (OCR) measured during static head tilts, and dynamic OCR quantified during off-vertical axis rotation (i.e vertical component of eye movement during off-vertical axis rotation, see (Beraneck et al., 2012)). Similarly to its effect on canal function, IDPN leads to a significant decrease of both the static OCR (Figure 1.C) and dynamic OCR (Figure 1.D) gain for the 3 highest doses (static OCR F(4, 31)=32.15 p<0.0001, [Control] vs [24], [Control] vs [32] and [Control] vs [40] p<0.001; Dyn OCR F(4, 31)=30.3 p<0.0001, [Control] vs [24], [Control] vs [32], and [Control] vs [40] p<0.001), and an intermediate loss for the [24] group (static OCR [16] vs [24], p=0.0021; [24] vs [32], p=0.023; Dyn OCR [16] vs [24], p<0.001; [24] vs [32], p=0.128).

Taken together, these results reveal the dose-dependent effect of IDPN on the canal- and otolith-specific VORs in mice.

### Dose-dependent effects of the treatment on the vestibular hair cells

To quantify the effect of the IDPN treatment on the vestibular HC, the vestibular organs were removed and fixed in paraformaldehyde (4%) after VOR measurements at D21. Immunochemistry was used to identify type I and type II HC with type-specific markers (respectively osteopontin (SPP1) and calretinin (Calre)(Borrajo et al., 2024)). The total number of HC, independently of their identity, was quantified using the pan-hair cell marker Myo7a. Figure 2.A shows typical examples of confocal (Zeiss LSM 880) immunostaining data obtained from [Control], [24] and [40] groups.

**Figure 2.**
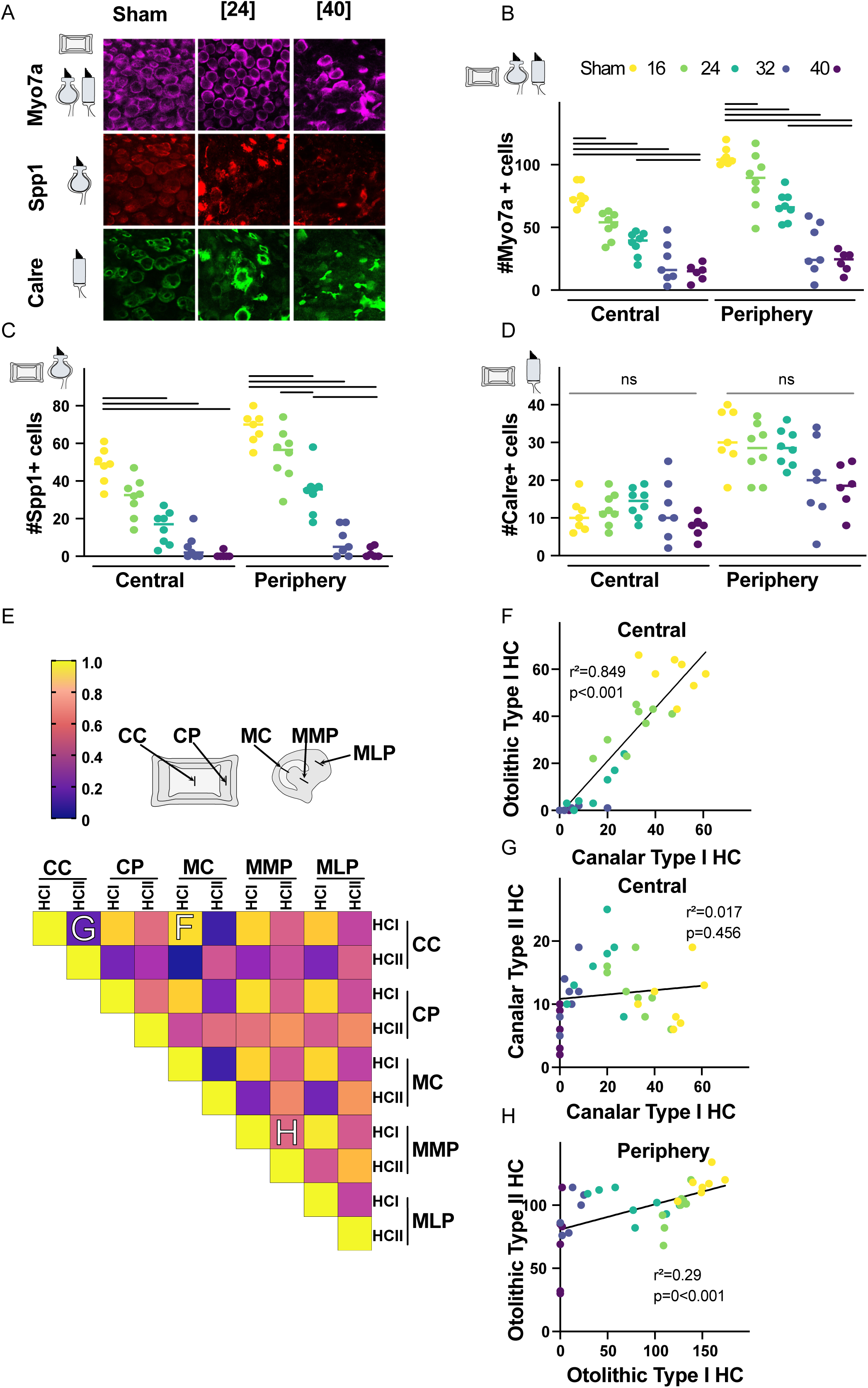
Dose-dependent ototoxic effect on vestibular hair cells. A) Immuno-labelling of all hair cells (Myo7a+), type I HC (Spp1+) and type II HC (Calre+) in the central ampulla of the horizontal canal in SHAM [0] (left panels), and mice injected with [24] (middle panels)] or [40] mmol/kg of IDPN. Samples were obtained at D21 following IDPN injection. (B, C, D) Cell count of hair cells in the central and peripheral horizontal ampulla for all IDPN doses, with (B) Myo7a+ marker (all HC) (C) Spp1+ marker (type I specific marker), (D) Calre+ marker (type II specific). Sample size Sham (n=7), [16] (n=8), [24] (n=8), [32] (n=7) and [40] (n=6). E) Correlation matrix of the r Pearson coefficient of the number of HCI and HCII for the n=36 mice in different regions of ampulla and macula (Grey scheme; C.C: crista central, C.P: crista periphery, M.C: macula central, M.M.P: macula medial periphery, M.L.P: macula lateral periphery). The color code representing the values of the coefficient are in the top left panel. F) Relation between number of type I HC in the macula and the number of type I HC in the ampula in the central region of vestibular organs of individuals. The linear fit is represented. G-H) Relation between number of type II HC in the ampulla and the number of type I HC in the ampulla in the central (G) or peripheral (H) regions of individuals. The linear fit is represented.

The results of HC quantification are plotted in Figure 2.B, C and D, respectively for Myo7a (all HC), SPP1 (type I-specific) and Calre (type II-specific) for both central and peripheral part of the horizontal crista (see Supplementary Figure 2 for otolith cell counting). IDPN exposure leads to a significant decrease in the number of Myo7a-labelled HC in the central part for all doses and in the peripheral region for doses above [16] (see Figure 2.B). The [24] treatment reduces by ∼50% Myo7a+ labelled cells compared to the Control group (Central: [Control] vs [24], p<0.0001; Peripheral: [Control] vs [24], p<0.0001). The two highest doses, though not different from each other (Central: [32] vs [40] and Peripheral: [32] vs [40] p>0.99), lead to a further 25% decrease in the number of Myo7a+ cells compared to the [24] group (Central: [24] vs [40] p=0.002; Peripheral: [24] vs [40] p<0.0001).

The decrease in the number of vestibular type I HC labelled by SPP1 is shown in Figure 2.C with a similar trend than the Myo7a labelling: loss is significant for all doses > [16] in the central and the peripheral regions (Central: [Control] vs [24] and Peripheral: [control] vs [24], p<0.0001). In contrast, IDPN did not lead to any change in the number of type II HC (Calre labelled cells) in the central part of the crista, and to a small and non-significant decrease limited to the highest dose of 40mmol/kg in the peripheral region (Central: [Control] vs [40] p>0.99; Peripheral: [Control] vs [40] p=0.8066).

To determine how the HC loss compares in both organs for all individuals, we compared Type I and Type II HC in all regions (Figure 2.E, top panel) by plotting a heat map based on the Pearson correlation matrix (Figure 2.E, bottom panel). While the correlation between type I HC remaining after IDPN is high in all regions, the correlation of Type II HC with type I HC is low. Figure 2.F, 2.G and 2.H illustrate the different sorts of correlation found between the HC identity and regions. The high correlation between the remaining Type I HC in the otolith and canal organs (Central: r²=0.849, p<0.001; Figure 2.F) shows that the IDPN treatment leads to a highly correlated loss in both structures for all doses. However, there is no correlation between the loss of Type I and Type II HC in the central regions of either organ (Canal: r²=0.017, p=0.456; Figure 2.G) and a small yet significant correlation in the peripheral regions (Otolithic r²=0.29, p<0.001; Figure 2.H) mostly driven by the small decrease in type II HC at the highest dose. This analysis also shows that the ototoxicity similarly affects canals and otoliths. Overall, the effect of the IDPN treatment leads to a largely specific dose-dependent loss of type I HC, with a comparable effect in central and peripheral regions of the ampulla and utricule.

Organ-specific structure-function relationship.

To investigate the relation between type I HC population of the different organs and the related vestibular function, we first compared the normalized individual gains of the vestibulo-ocular reflexes and the normalized number of type I (SPP1-labelled cells) HC in the central region of the canals (Figure 3.A and 3.B) and otoliths (Figure 3.C and 3.D). Normalization values were obtained using the mean of the control group (yellow dots) as maximal function (100%).

**Figure 3:**
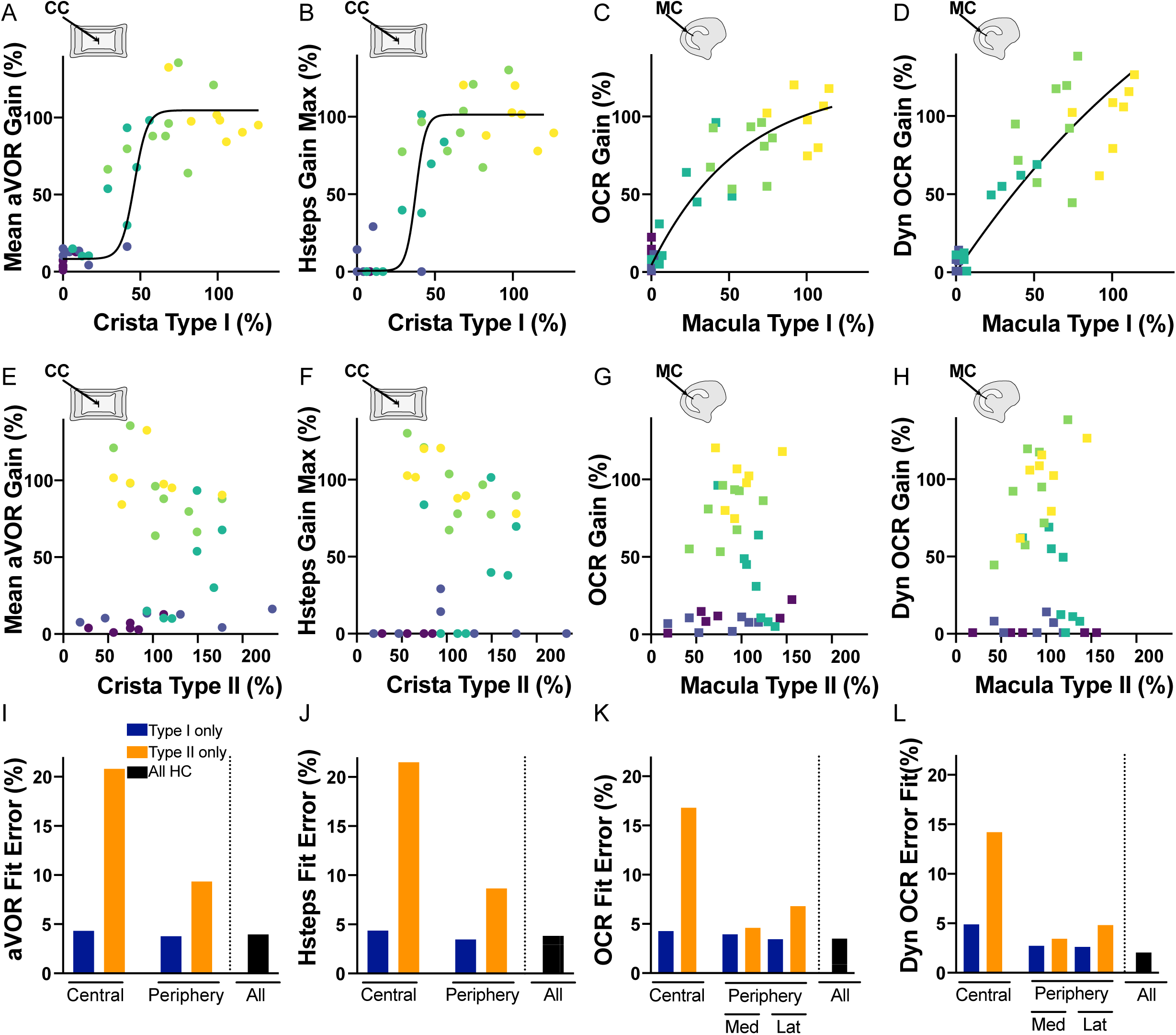
Relation between hair cells subtype and vestibulo-ocular reflexes. A,B,C,D) Normalized Mean aVOR (A), Hsteps (B), OCR (C) OCR Dyn (D), as a function of the normalized number of Type I hair cells in the central part of the ampulla (A, B) or the central part of the macula (C, D). The non-linear 2D fit is represented. E,F,G,H) Normalized Mean aVOR (E), Hsteps (F), OCR (G) OCR Dyn (H), as a function of the normalized number of Type II hair cells in the central part of the ampulla (E,F) or the central part of the macula (G,H). The non-linear 2D fit is represented. I,J,K,L) Error Fit (%) of the 2D sigmoid fit for the Mean aVOR (I), Hsteps (J), OCR (K) OCR Dyn (L) in the central region and peripheral regions of the ampulla (I, J) or the macula (K, L) for type I HC, type II HC and All Types (Type I + Type II).

The relationship between the number of type I HC and the vestibular functions was approximated by interpolating the data with 2D sigmoidal curves accounting for both the vertical (measured function values) and horizontal (HC staining data) variability. Figure 3.A-D (canals, Figure 3.A and B; otoliths Figure 3.C and D) shows that the non-linearity of the HC-function relationship is particularly evident for canal tests. Figure 3.E-H illustrate the absence of correlation between the gains measured and the number of type II HC counted in the central regions of both vestibular organs (canals, Figure 3.E-F; otoliths Figure 3.G-H). Since IDPN treatment differentially affects specific regions and cell types (Figure 2), we calculated the different Fit Error (%) of the sigmoids to determine which region/cell type better relates to the various VOR tests. This parameter is plotted for each test, and for the central and peripheral regions of each organ, considering either type I (SPP1 marker), type II (Calre marker) or both HC (type I + type II) (aVOR, Figure 3.I; HSteps Figure 3.J; OCR Figure 3.K; Dyn OCR Figure 3.L). In all instances, models based on type I HC always better approximate the function than models based on type II HC (i.e make less error; compare blue and yellow plots). Models based on type II HC in the periphery perform better than models based on type II HC in central parts, which probably relates to the partial and small dose-dependent effect found for the highest concentrations of IDPN (e.g. Fig 2.D, G, H). Finally, using a model summing central and peripheral type I and type II HC led to a similar (3.I-K) or smaller (3.L) error and thus better represented all the different VOR tests (compare dark and colored plots in panels 3.I-L).

Overall, these results demonstrate that the responses to these different VOR tests strongly depend on the integrity and proportion of type I HC. In the instances where all type I HC are lost, the presence of type II HC alone does not allow to preserve the function. These targeted analyses also suggested that the different VOR tests have differential dependency on the amount of HC preserved. We therefore decided to investigate which quantity of total HC is necessary for the maintenance of minimal or normal VOR function.

### Thresholds of minimal and normal population-coding in canal and otolith functions

To determine these thresholds, we first grouped the mice based on their loss of type I HC observed in the different organs. As shown in Figure 4.A, there is a large intra-group variability and inter-group overlap in the amount of HCI left in all regions, leading to an apparent continuum of type I HC integrity that ranges from the control animals (yellow) to the highest dose of the treatment ([40], dark purple). However, a cluster analysis on this dataset revealed 3 distinct categories of animals characterized by high (green), intermediate (orange), or low (pink) type I HC population integrity (Figure 4.B). We then validated that Myo7a is a pan-hair cell marker which in both canals (Figure 4.C) and otoliths (Figure 4.D) closely matches the number of both type I and type II HC. The high correlation between the addition of type I + type II and myo7a-labelled cells (Canals r²=0.99, p<0.0001 Figure 4.C; Otoliths r²=0.979, p<0.0001, Figure 4.D) is maintained for all 3 clusters, demonstrating that despite the ototoxic alteration of the HC, this marker remains an accurate index of total HC integrity in all our experimental conditions.

**Figure 4:**
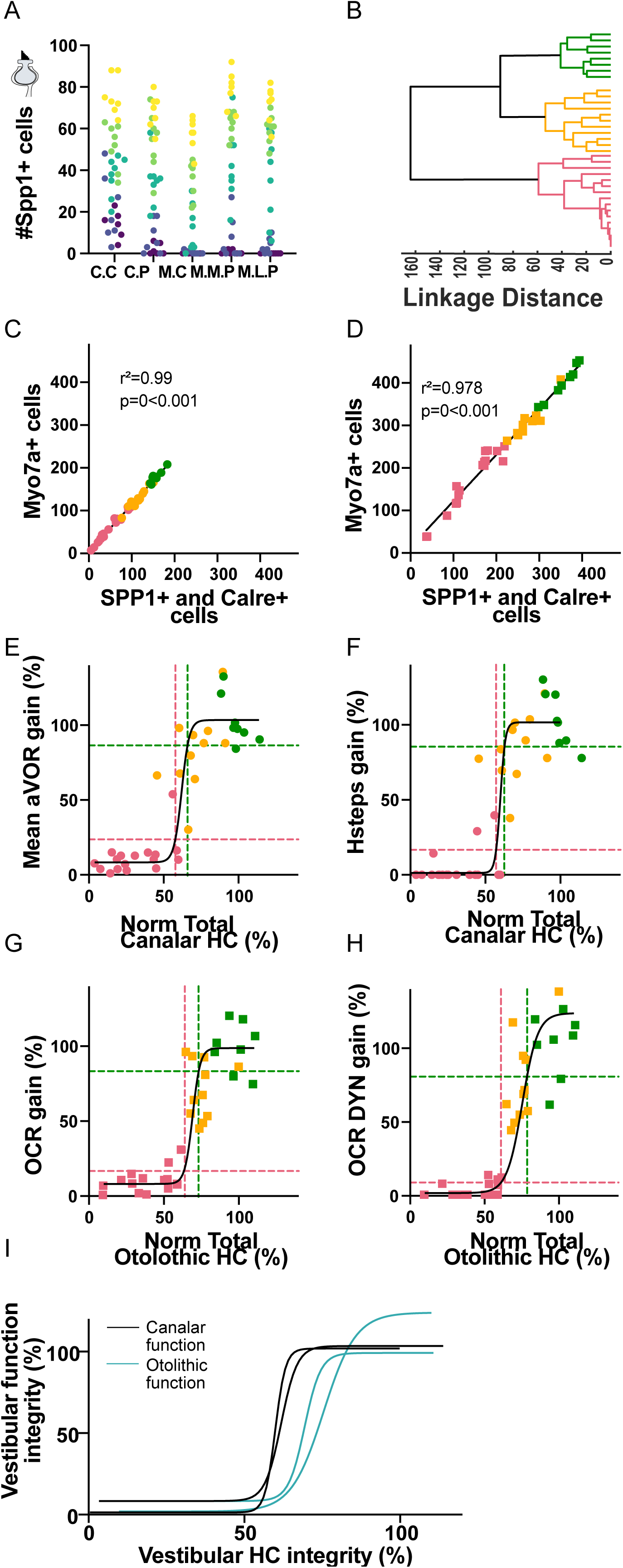
proportion of hair cells to maintain minimal and normal vestibular responses. A) Number of type I hair cells counted in all regions of ampulla and macula (C.C: crista central, C.P: crista periphery, M.C: macula central, M.M.P: macula medial periphery, M.L.P: macula lateral periphery). B) Clustering dendrogram of the individuals based on their number of type I hair cells in the different regions shown in (A). C, D) Number of Myo7a-labelled hair cells as a function of the additive number of Spp1- and Calre-labelled hair cells in the ampulla (C) and macula (D). E, F) Normalized Mean aVOR gain (E) and HSteps gain max (F) as a function of the normalized number of Myo7a-labelled hair cells in all regions of the ampulla. G, H) Normalized static OCR gain (G) and Dynamical OCR gain (H) as a function of the normalized number of Myo7a-labelled hair cells in all regions of the macula. In panel E-G the lowest (highest) threshold is obtained as the value of the sigmoidal fit, black line, for value of mean + (-) standard deviation of the pink (green) cluster group. I) Superimposed 2D Non-linear fits of the vestibular hair cells integrity (%) as a function of the percentage of canal (black) and otolithic function (blue).

In Figure 4.E-H, a comprehensive analysis of the global structure-function relationship encompassing all the hair cells was conducted by applying to the total hair cell (HC) residual number the same sigmoidal fitting procedure employed for the specific HC types and locations (Figure 3). For each test, we defined the minimal function thresholds as the “mean+SD” gain of the low response cluster (pink dotted lines), while the normal vestibular function threshold was defined as the “*mean-SD*” gain of the high response cluster (green dotted lines). Using this methodology, normal thresholds represented >80% of function, while minimal threshold represents < 20% of function (ordinate of the horizontal dotted line on panels 4E-H).

The corresponding HC thresholds were then measured as the intercept between the function thresholds and the sigmoid curve. The quantity of HC to preserve minimal horizontal canal function was calculated at 57.8% for aVOR (Figure 4.E) and 57.03% for Hsteps (Figure 4.F), while the threshold for normal functions were calculated at 65.9% and 62.5%, respectively. Minimal HC thresholds for otolithic-dependent responses were slightly higher with lower bounds calculated at 63.5% and 60.9% respectively for static OCR (Figure 4.G) and Dynamic OCR (Figure 4.H). The HC thresholds for normal otolithic function were calculated at 73.1% and 78.5%, respectively, slightly higher than for canal function. This analysis suggests that canal-dependent function tested in our protocol remains in normal range when roughly 2/3 of HC are preserved, but requires about 1/2 of HC to preserve minimal function. On the other hand, otolith-dependent function tested in our protocol requires approximatively 2/3 of HC to maintain minimal function and ¾ of HC to remain in normal range,

To further highlight the relation between the total number of hair cells in ampulla and macula and organ-specific function, the non-linear fits of the normalized vestibular function presented in panels 4E-H are superimposed on Figure 4I. This summary scheme illustrates that despite the experimental variability in measurements, inter-individual variability, and differences in the parameters tested (static vs dynamic; long-lasting vs instantaneous responses), all vestibulo-ocular reflexes show a rather comparable dependency on the HC population integrity necessary to drive the eye movements, with no response observed when 50% of hair cell are lost, intermediate response when HC population is in range 50-80%, and normal function for both organs when HC integrity is >80%.

## DISCUSSION

This study aimed to determine the minimal proportion of surviving vestibular HC necessary to maintain normal vestibular function, specifically assessed via the vestibulo-ocular reflex. Our experimentation involved exposing mice to increasing doses of an ototoxic compound, IDPN (3,3’-Iminodipropionitrile), and revealed that vestibular function declines progressively, until vestibular responses are completely abolished. By combining immunohistochemistry with video-oculography in individuals, we found that VOR performance remains physiologically normal as long as at least 80% of type I HCs are preserved. We further found that no VOR response was detectable when fewer than 50% of type I HCs were preserved. To our knowledge, these thresholds represent the first quantitative benchmarks linking cellular survival to a quantifiable behavioral output of vestibular function, with relevance across species and potential translatability to clinical applications in humans.

### A single injection of IDPN at different doses is able to induce graded HC loss

We recently demonstrated (Schenberg et al., 2023) that IDPN (3,3’-Iminodipropionitrile) administering at low doses in drinking water could cause a progressive and reversible alteration of type I HCs resulting in VOR dysfunction followed by recovery. This subchronic exposure protocol revealed that type I HCs are essential for maintaining VOR function. However, that previous approach did not allow for a quantitative assessment of the level of vestibular HC integrity required for proper VOR function.

In the present work, we used single IDPN injections at various concentration to induce a permanent loss of HCs. Our findings corroborate prior studies (Llorens et al., 1993 p.93; Soler-Martin et al., 2007; Martins-Lopes et al., 2019; Zeng et al., 2020; Maroto et al., 2021), confirming that IDPN induces a dose-dependent and graded vestibular HCs loss, as evidenced by the quantification performed with the pan-HC marker Myo7a.

We validated the dynamics of functional loss in animals after a single injection at 24mM (n=3) and 40mM (n=2) (data not shown). VOR decline was partial within the first 4 days post-injection, reaching complete loss after one week. To confirm the stability of our measurements, all animals of this study were tested at two- and three-weeks post-injection, confirming that VOR responses remained stable between these time points (Supplementary Figure 1). While other ototoxic drugs such as cisplatin have been shown to induce a plateau in hair cell loss even at high doses (Ding et al., 2018), IDPN represents a cost-effective and practical pharmacological tool for inducing a wide range of permanent vestibular HC loss in mice, from minimal to complete (Llorens et al., 1993; Soler-Martín et al., 2007; Maroto et al., 2021).

### Quantification of the differential HC loss in subregions of ampulla and macula

It has long been recognized that the HCs found in the different regions of the vestibular organs differ by several characteristics (Goldberg, 2000; Eatock and Songer, 2011). Type I HCs and their associated calyx-shaped synapse were originally thought to be more numerous in the apex/center of the cristae ampullaris and in the central striola region of the maculae of the otolith organs (Eatock and Songer, 2011), with type II HCs more prevalent in peripheral and non-striolar zones of the different vestibular organs, which are also characterized by a higher density of regular afferents (Goldberg, 2000). In our study, although we observed a higher number of type I HC compared to type II HC in the central regions, peripheral regions still showed a substantial number of type I HC.

Here, we conducted organ-specific vestibular tests in order to specifically assess the stabilizing eye movements that depend on the semicircular canals or otolith organs. As illustrated in Figure 2E, there was, however, a strong correlation between the HC loss observed in both set of organs, as well as between the different subregions. Type I HCs were lost in a dose-dependent fashion, and the loss was found to affect all organs and subregions uniformly. As such, our results do not allow us to distinguish between the specific roles played by the type I HCs located in central versus peripheral areas of the organs. Similarly, type II HCs were mostly unaffected by IDPN, except at the highest dose tested. Indeed, in some individuals of the [32] and [40] groups, there was a moderate, non-significant, loss in the periphery of the organs. However, we note that these are the areas where the type II HCs are more abundant (Figure 2D) and therefore, where the subtler effects of IDPN would be most readily detected by our quantification. Consequently, the main point that this dataset allows us to investigate lies in the role of type I and type II HCs in the encoding of head movements. As previously mentioned, type I and II vestibular HCs are hypothesized to form complementary sensory channels, enabling the brain to encode distinct features of head position and motion in space.

The type I HCs and their associated calyceal synapse, with both quantal and non-quantal modes of transmission, appear particularly well-suited for rapid information processing, likely essential for short-latency reflexive vestibular pathways. Thus, the encoding of rapid head movements with high-frequency content, whether generated during natural head movements (Carriot et al., 2017), bone-conducted vibrations (Curthoys et al., 2017), or jerk stimuli used to evoke vestibular sensory evoked potential (VsEPs; Jones et al., 2011), is thought to depend predominantly on type I HCs. In contrast, type II HCs with their button-type synaptic endings, are hypothesized to be better suited for encoding low dynamic, tonic and/or sustained stimuli, such as static head tilts. However, it is well recognized that most head movements simultaneously activate all type of HCs (Eatock, 2018), and that some afferents may receive inputs from neighboring type I and type II HC (Lysakowski and Goldberg, 1997; Goldberg, 2000).

In our study, the loss of type I, but not type II, HCs led to a marked reduction in responses across all VOR tests. The presence of type I HCs is therefore shown to be critical for encoding all types of head movements, ranging from static head tilt to horizontal rotation with low to moderate dynamics. During static head tilt, head position relative to gravity is encoded as a change in the tonic firing rate of afferents (Goldberg and Fernandez, 1975), with better discrimination from regular than irregular afferents (Jamali et al., 2016). While this kind of stimulus with slow, sustained movement is usually thought to depend on the tonic neural elements (i.e type II HC and regular afferents), our findings indicate that the type II HCs, located in both the central and peripheral parts of the otoliths and largely unaffected by the treatment, are insufficient on their own to mediate otolith-driven eye movements.

Overall, these observations confirm the critical role of type I HCs in the encoding of VOR, demonstrate that type II HCs alone are insufficient to drive compensatory eye movements, and suggest that both hair cell types are required to appropriately drive the discharge of vestibular afferents.

### Correlation of function with HC integrity indicate minimal and normal thresholds

By correlating structural and functional measures in individual subjects, we found that restoring at least 60% of overall HC integrity is likely required to support minimal vestibular function, while approximately 80% appears necessary to achieve near-normal responses. Importantly, these thresholds reflect the total HC population and not specifically type I cells, although our findings suggest that type I HCs are especially critical for robust responses. In our protocol, VOR performance reflects contributions from both type II HC and the surviving population of type I hair cells, making it difficult to disentangle the specific role of each subtype in driving compensatory eye movements. When considering type I cells in isolation, thresholds for minimal and near-normal responses would be lower—around 40% and 60%, respectively. As such, our current estimates are conservative and may overstate the actual proportion of HC required. Further studies will be needed to determine whether similar functional outcomes can be achieved with a smaller number of HCs, provided that a greater proportion of type I HCs is preserved.

In addition, by using tests specifically designed to differentiate canal-versus otolith-driven eye movements, we observed differences in the functional thresholds derived from both sets of organs (Figure 4). These differences could be attributed to experimental limitations. For instance, the number of measurements acquired for each test differed, with fewer trials conducted for the otolithic tests compared to the canal-based ones, potentially affecting the resolution of the functional assessments. Moreover, the higher number of hair cells present in the otolithic organs relative to the semicircular canals could introduce a quantification bias.

Alternatively, this difference may reflect distinct population-coding strategies between the semicircular canals and the otolithic organs. In the ampullae, all HCs are activated by the same directional acceleration—i.e., based on head movements relative to the orientation of the main axis of the semicircular canal. In contrast, the maculae exhibit a more heterogeneous organization, with hair cells oriented along varying axes across different subregions. As a result, encoding a specific directional stimulus in the otolith organs may require a larger and more spatially distributed population of hair cells to achieve sufficient signal resolution (Simon et al., 2021).

### Restorative thresholds for vestibular recovery in gene therapy

Recent advancements in gene therapy, including the use of adeno-associated virus vectors (Isgrig et al., 2017), antisense oligonucleotides (Lentz et al., 2013) or even lentiviral vectors (Schott et al., 2023), have shown promise in restoring vestibular HCs and their associated functions. Our findings offer valuable quantitative benchmarks to guide the development and evaluation of such therapeutic strategies.

The minimal and near-normal thresholds of vestibular function identified at 60 and 80%, respectively, align with results from recent gene therapy investigations employing comparable methodologies. For instance, (Schlecker et al., 2011) demonstrated that balance impairments induced by IDPN exposure could be reversed following adenoviral-mediated gene delivery, which resulted in near-complete restoration of utricular HCs, including a proportion of type I HCs. Similarly, in a study by (Jáuregui et al., 2024), retention of approximately 50% of type I HCs after diphtheria toxin–induced damage was associated with significant preservation of vestibular function. When diphtheria toxin was also utilized to selectively target HCs, the regeneration of both type I and II otolithic HCs, ensured via Wnt pathway activation, achieved up to 58% of the original HC population. However, the majority of the regenerated cells exhibited morphological and molecular features characteristic of type II HCs, with relatively few type I HCs restored. As a consequence, only partial functional recovery was observed, with modest improvements limited to the higher frequencies of the rotational VOR. In contrast, regeneration rates below 50%—with utricular and canalar HC counts restored to only 21% and 14%, respectively—were insufficient to yield any measurable improvement in vestibulo-ocular reflexes (Lahlou et al., 2024b). These findings, taken together with our data, suggest that effective gene- or pharmacologically-based therapies for vestibular dysfunction should aim to restore at least 50–60% of the vestibular HC population to produce meaningful functional recovery. Notably, achieving higher proportions of regenerated type I HCs may be particularly important to support full restoration of reflexive vestibular responses.

A major limitation of current therapeutic strategies lies in their predominant emphasis on promoting the recovery of type II HC, resulting in either absent or suboptimal recovery of vestibular function (Jáuregui et al., 2024; Lahlou et al., 2024b). In neonatal mice, satellite glial cells facilitate the regeneration of both type I and II HCs, enabling more complete restoration of vestibular HCs (Wang et al., 2015). In contrast, spontaneous regeneration of HCs in adult mice is limited and occurs primarily in response to injury. Notably, the newly regenerated vestibular cells exhibit a limited range of HC subtypes as regenerated cells have not been reported to include type I-like HCs (González-Garrido et al., 2021). Most newly formed cells display features characteristic of type II HCs, while others remain in immature or undifferentiated states (Kawamoto et al., 2009; Hicks et al., 2020; González-Garrido et al., 2021). This bias toward type II HC regeneration is also reflected in the preferential re-establishment of bouton-type afferent synapses, as opposed to calyx afferents that are associated with type I HCs (Zakir and Dickman, 2006). Despite this, pharmacological interventions have shown that both regular and irregular afferent activities can be reinstated following HC loss (Lahlou et al., 2024). Together, these findings underscore a critical need for future therapeutic approaches to not only enhance HC regeneration but also ensure the recovery and functional integration of both type I and type II vestibular HCs. This dual focus will be essential to achieve optimal and physiologically meaningful restoration of vestibular function.

### Perspectives and conclusion

In our study, vestibular function was assessed through measurements of the vestibulo-ocular reflex (VOR), a well-characterized and quantifiable output of vestibular processing. However, the vestibular system contributes to a broad array of reflexive and cognitive functions beyond the VOR, including vestibulo-spinal reflexes, postural control, and higher-order processes such as spatial orientation and navigation (Cullen and Taube, 2017). Whether the functional thresholds we identified—approximately 60% and 80% of HC integrity for minimal and near-normal VOR responses, respectively—are generalizable to these other vestibular modalities remains an open question. For example, vestibulo-spinal reflexes, which underlie balance and gait stability, may exhibit different sensitivity to hair cell loss, potentially requiring distinct levels of cellular preservation for functional maintenance. Similarly, cognitive processes that rely on vestibular input but integrate multisensory information may show greater resilience to partial vestibular damage due to compensatory mechanisms. Moreover, it is important to consider that vestibular functional preservation may not be determined solely by the quantity of surviving hair cells. The integrity of synaptic connections with vestibular afferents and the plasticity of central pathways, particularly within the brainstem, likely play key roles in shaping residual function and recovery potential.

In summary, our findings emphasize the pivotal role of type I HCs in maintaining vestibular function and validate IDPN-induced HC loss as a robust model for investigating vestibular decline, especially in the context of aging. Furthermore, the quantitative thresholds identified in this study offer valuable benchmarks for guiding future regenerative therapies aimed at restoring vestibular function, and underscore the need for comprehensive assessments that extend beyond the VOR to fully capture the complexity of vestibular system recovery.

## Figure legends

**Supplementary Figure 1:** Mean aVOR gain (A), HStep gain (B), static OCR gain (C), Dynamic OCR gain (D) of all (n=36) mice at D14 and D21 after injection.

**Supplementary Figure 2:** (A, B, C) Cell count of hair cells in the central and peripheral utricule for all IDPN doses, with (A) Myo7a+ marker (all HC) (B) Spp1+ marker (type I specific marker), (C) Calre+ marker (type II specific). Sample size Sham (n=7), [16] (n=8), [24] (n=8), [32] (n=7) and [40] (n=6).

## MATERIELS ET METHODS

### ANIMALS

A total of 36 male and female C57/BL6J mice, age 6–10 weeks, was used in this study. Mice were kept in standard lighting and housing conditions. Animals were used in accordance with the European Communities Council Directive 2010/63/EU. All efforts were made to minimize suffering and reduce the number of animals included in the study. All procedures were approved by the ethical committee for animal research of the Université Paris Cité.

### HEAD POST IMPLANTATION

Head post implantation surgery was performed as previously described(França de Barros et al., 2020). Mice were anesthetized with isoflurane, and a small longitudinal incision was made on their shaved head to expose the skull after a local injection of lidocaine hydrochloride (2%; 2 mg/Kg). A custom-built headpost 3×3×5mm; poly lactic acid) was first cemented (C&B Metabond; Parkell Inc., Edgewood, NY) and the sides were then covered with resin (Heraeus) for protection. Mice were placed under red light post-surgery until their full recovery and were monitored for the following 48h.

### OTOTOXIC EXPOSURE

Mice were treated with a single IP injection of IDPN. The animals were divided into 5 groups that received different doses of IDPN: 16 mmol/kg (n=8), 24 mmol/kg (n=8), 32 mmol/kg (n=7), 40 (n=6) mmol/kg and the control group (n=7) that received an injection of NaCl. Vestibular function was tested 2 (D14) and 3 weeks (D21) after the injection. As the longer time after exposure led to no further vestibular function deprivation (mix-model statistical test, non-significant differences between D14 and D21 for each test (aVOR, Hsteps, OCR and OCR Dynamic, see supplementary Figure 1), D14 and D21 functional data were pooled.

### VIDEO OCULOGRAPHY RECORDING SESSIONS

Eye movements were recorded with a non-invasive video-oculography system (ETL 200 Iscan, Acquisition rate 120 Hz) following procedure previously described (Stahl, 2004). In brief, mice were positioned in a Plexigas tube and head-fixed with a nose-down angle of 30° to align the horizontal semi-circular canals with the yaw plane. The tube was fixed on a rotating platform on top of an extended rig with a servo-controlled motor. Eye and Image/head position signals were sampled at 1 kHz, digitally recorded (CED power1401 MkII) with Spike 2 software and analyzed off-line in Matlab (Matlab, The MathWorks, Natick, MA, USA; RRID: SCR:001622) programming environment. Recording sessions were performed in a temperature-controlled room (21-24°) and lasted up to 45 min.

### VESTIBULAR STIMULATION

Canal (VORd and Hsteps) and utricular-specific (OCR and OCR Dynamic) tests were performed in complete darkness, with the mouse surrounded by an opaque dome or box. Pilocarpine 2% was placed on the eye of the mouse before the test to prevent excessive pupil dilatation.

Sinusoidal angular rotations around the vertical axis were performed to record the horizontal angular vestibulo-ocular reflex (aVOR), at different frequencies: 0.2, 0.5, 0.8; 1 and 2Hz at a peak velocity of 30°/s. At least 10 cycles were analysed per frequency and the compensatory eye movements were quantified by calculating the gain (ratio between the eye velocity and table velocity) and the phase (normalized latency between the eye and the table velocities) (Carcaud et al., 2017).

HSteps test were performed using an abrupt acceleration followed by a sudden stop after three 360° rotations with a constant-velocity of 50°/s on the yaw plane (acceleration 250°/s²). The hsteps gain was calculated as the ratio between the peak velocity of the slow phases at onset and offset of rotations, and the rotation velocity.

Static Counter Roll (OCR) tests were performed first by measuring the vertical pupil position according to the head tilt angle at 0° in the horizontal plane. The table was then tilted from left to right in incremental steps of 10° (from 0 to 40deg), with static periods of at least 10s between oscillations (Fig1.D) to record stable eye position. The OCR gain corresponds to the slope of the linear regression of the vertical eye angle and the head tilt angles^,31^.

Dynamic OCR tests were performed with the vestibular turntable tilted with a 17° off-axis angle (Beraneck et al., 2012; Idoux et al., 2018; Simon et al., 2021). 50°/s continuous stimulations were performed in a counter-clockwise and then clockwise direction. The Dynamic OCR gain corresponds to the dynamic, or vertical component of the compensatory eye movement divided by the maximal differential tilt angle (i.e 34°).

### IMMUNOLABELLING OF THE HAIR CELLS IN THE HORIZONTAL SEMI-CIRCULAR CANALS AND THE UTRICULE

The inner ear of all mice tested with vestibular function tests (n=36) were then used to perform immunofluorescence analysis on hair cells in the vestibular endorgans. Mice were anaesthetized with an overdose of intraperitoneal injection of ketamine hydrochloride (10%)/xylazine (5%) and decapitated. The histology was done following previously published protocol of Maroto and colleagues (Maroto et al., 2021). The vestibular epithelia were dissected and fixed for 1h in a 4% solution of paraformaldehyde (PFA). PFA was washed twice with phosphate-buffered saline (PBS) and the samples were placed in a cryoprotective solution at 4° for 2h for effective embedding and then stored at -20°. Before the immunochemistry, samples (one horizontal canal and one utricule) were put at room temperature and rinsed twice in PBS. While under slow agitation, the samples were incubated twice, first for 1h with 4% Triton X-100 (Sigma Aldrich) in PBS to permeabilise the tissue and a second time for 1h in 0.5% Triton X-100 1% fish gelatin (CAS #9000-70-8, Sigma-Aldrich) in PBS to block non-specific binding sites. The incubation with the primary antibodies was then performed in 0.1% Triton X-100, 1% fish gelatin in PBS at 4° for 24h. After rinsing, the secondary antibodies were incubated in the same conditions. The secondary antibodies were rinsed and the vestibular epithelia were mounted on slides with fluoromount (F4680, Sigma-Aldrich) and were visualised with a Zeiss LSM880 confocal microscope (with an objective of 63x NA:1;4). To properly analyse the whole vestibular epithelium, Z-stacks of 0.5 μm were obtained and observed with ImageJ (National Institute of Mental Health, Bethesda, Malyland, USA). The primary antibodies used were rabbit anti-Myosin VIIa (Myo7a) (Proteus Biosciences, #25-6790, 1:400), guinea pig anti calretinin (Synaptic Systems #214-104, 1:500) and goat anti-osteopontin (SPP1) (R&D Systems #AF08, 1:200). Their respective secondary antibodies were Dylight 405 donkey anti-rabbit ifG H+L (Jackson Immuno Research #711-475-152, Alexa Fluor 488 donkey anti-guinea-pig IgG H+L (Jackson ImmunoResearch #706-605-148) and Alexa Fluor 555 donkey anti-goat IgG H+L (Invitrogen #A21432).

The global number of hair cells was assessed with the cytoplasmic labelling of the anti-Myo7a antibody(Hasson et al., 1997). Type I hair cells were labelled using anti-SPP1, found in the neck of the type I HC (McInturff et al., 2018). Type II hair cells were distinguished by the colocalization of Myo7a and calretinin (Dechesne et al., 1991). A thorough evaluation of the specificity of these markers in the adult rat has been recently published (Borrajo et al., 2024).

## Acknowledgment

This study contributes to the IdEx Université de Paris ANR-18-IDEX-0001. This work has benefited from the support and expertise of the animal facility of BioMedTech Facilities at Université Paris Cité ((https://biomedicale.u-paris.fr/biomedtech-facilities/) INSERM US36 | CNRS UAR2009 | Université Paris Cité). The confocal microscopy studies were performed at the Centres Científics i Tecnològics de la Universitat de Barcelona (CCiTUB). We thank Dr. Benjamin Torrejon for technical assistance and Dr. Alberto Maroto for preliminary work for this study.

## Author contribution

Conceptualization: JL; MB. Methodology: LS; AP; FS; MT; JL; MB. Formal analysis: LS; AP; CD; MT; JL; MB. Investigation: LS; AP; CD; MB. Writing original manuscript: LS; MB. Writing review and editing: LS;; FS; MT; JL; MB. Visualization: LS; AP; MT; JL; MB. Supervision: LS; FS; MT; JL; MB. Project administration: JL; MB. Funding acquisition: JL; MB.

## STATISTICAL ANALYSIS

To test the main effect of the between-individual factor ‘Dose’ (0, 16, 24, 32 or 40 mmol/kg) on Hsteps, OCR and DYN OCR, as well as the effect of IDPN exposure on the hair cells, a one-way ANOVA was used. For the aVOR a two-ways ANOVA was used to be able to evaluate also the main effect of the within-individual independent factor ‘Frequency’ (0.2, 0.5, 0.8, 1 and 2Hz) and its interaction with the Dose. The difference between D14 and D21 vestibular reflex tests was reported with a mixed model analysis. The significance threshold was set at p<0.05 and Tukey multiple comparison test was performed if the main effect, or an interaction, was reported significant.

To describe the relationship between cell staining and the behavioural measures, 2D non-linear (sigmoidal) models were used to fit the datasets. Given the presence of noise on both types of measurement (function and cell counting data), the fitting required orthogonal distance regression and the fit was optimized for each data set.

## BIBLIOGRAPHY

Agrawal Y, Ward BK, Minor LB (2013) Vestibular dysfunction: Prevalence, impact and need for targeted treatment Chabbert C, ed. VES 23:113–117.

Angelaki DE, Cullen KE (2008) Vestibular System: The Many Facets of a Multimodal Sense. Annu Rev Neurosci 31:125–150.

Anon (n.d.) Evolutionary and Developmental Biology Provide Insights Into the Regeneration of Organ of Corti Hair Cells - PubMed. Available at: https://pubmed.ncbi.nlm.nih.gov/30135646/ [Accessed April 10, 2025].

Beraneck M, Bojados M, Le Séac’h A, Jamon M, Vidal P-P (2012) Ontogeny of Mouse Vestibulo-Ocular Reflex Following Genetic or Environmental Alteration of Gravity Sensing Gilestro GF, ed. PLoS ONE 7:e40414.

Borrajo M, Sedano D, Palou A, Giménez-Esbrí V, Barrallo-Gimeno A, Llorens J (2024) Maturation of type I and type II rat vestibular hair cells in vivo and in vitro. Front Cell Dev Biol 12:1404894.

Carcaud J, França de Barros F, Idoux E, Eugène D, Reveret L, Moore LE, Vidal P-P, Beraneck M (2017) Long-Lasting Visuo-Vestibular Mismatch in Freely-Behaving Mice Reduces the Vestibulo-Ocular Reflex and Leads to Neural Changes in the Direct Vestibular Pathway. eNeuro 4:ENEURO.0290-16.2017.

Carriot J, Jamali M, Chacron MJ, Cullen KE (2017) The statistics of the vestibular input experienced during natural self-motion differ between rodents and primates: Natural vestibular input in rodents and monkeys. J Physiol 595:2751–2766.

Contini D, Holstein GR, Art JJ (2024) Simultaneous recordings from vestibular Type I hair cells and their calyceal afferents in mice. Front Neurol 15:1434026.

Cullen KE, Taube JS (2017) Our sense of direction: progress, controversies and challenges. Nature Neuroscience 20:1465.

Curthoys IS, MacDougall HG, Vidal P-P, de Waele C (2017) Sustained and Transient Vestibular Systems: A Physiological Basis for Interpreting Vestibular Function. Front Neurol 8 Available at: http://journal.frontiersin.org/article/10.3389/fneur.2017.00117/full.

Dechesne CJ, Winsky L, Kim HN, Goping G, Vu TD, Wenthold RJ, Jacobowitz DM (1991) Identification and ultrastructural localization of a calretinin-like calcium-binding protein (protein 10) in the guinea pig and rat inner ear. Brain Research 560:139–148.

Ding D, Jiang H, Zhang J, Xu X, Qi W, Shi H, Yin S, Salvi R (2018) Cisplatin-induced vestibular hair cell lesion-less damage at high doses. J Otol 13:115–121.

Eatock RA (2018) Specializations for Fast Signaling in the Amniote Vestibular Inner Ear. Integrative and Comparative Biology 58:341–350.

Eatock RA, Songer JE (2011) Vestibular Hair Cells and Afferents: Two Channels for Head Motion Signals. Annu Rev Neurosci 34:501–534.

Emptoz A, Michel V, Lelli A, Akil O, Boutet De Monvel J, Lahlou G, Meyer A, Dupont T, Nouaille S, Ey E, Franca De Barros F, Beraneck M, Dulon D, Hardelin J-P, Lustig L, Avan P, Petit C, Safieddine S (2017) Local gene therapy durably restores vestibular function in a mouse model of Usher syndrome type 1G. Proc Natl Acad Sci USA 114:9695–9700.

Fleihan T, Nader ME, Dickman JD (2024) Cisplatin vestibulotoxicity: a current review. Front Surg 11:1437468.

França de Barros F, Schenberg L, Tagliabue M, Beraneck M (2020) Long term visuo-vestibular mismatch in freely behaving mice differentially affects gaze stabilizing reflexes. Sci Rep 10:20018.

Goldberg JM (2000) Afferent diversity and the organization of central vestibular pathways. Exp Brain Res 130:277–297.

Goldberg JM, Fernandez C (1975) Responses Of Peripheral Vestibular Neurons To Angular And Linear Accelerations In The Squirrel Monkey. Acta Oto-Laryngologica 80:101–110.

Golub JS, Tong L, Ngyuen TB, Hume CR, Palmiter RD, Rubel EW, Stone JS (2012) Hair cell replacement in adult mouse utricles after targeted ablation of hair cells with diphtheria toxin. J Neurosci 32:15093–15105.

González-Garrido A, Pujol R, López-Ramírez O, Finkbeiner C, Eatock RA, Stone JS (2021) The Differentiation Status of Hair Cells That Regenerate Naturally in the Vestibular Inner Ear of the Adult Mouse. J Neurosci 41:7779–7796.

Halmagyi GM, Fattore CM, Curthoys IS, Wade S (1994) Gentamicin vestibulotoxicity. Otolaryngol Head Neck Surg 111:571–574.

Hasson T, Gillespie PG, Garcia JA, MacDonald RB, Zhao Y, Yee AG, Mooseker MS, Corey DP (1997) Unconventional Myosins in Inner-Ear Sensory Epithelia. Journal of Cell Biology 137:1287–1307.

Hicks KL, Wisner SR, Cox BC, Stone JS (2020) Atoh1 is required in supporting cells for regeneration of vestibular hair cells in adult mice. Hear Res 385:107838.

Huterer M, Cullen KE (2002) Vestibuloocular Reflex Dynamics During High-Frequency and High-Acceleration Rotations of the Head on Body in Rhesus Monkey. Journal of Neurophysiology 88:13–28.

Idoux E, Tagliabue M, Beraneck M (2018) No Gain No Pain: Relations Between Vestibulo-Ocular Reflexes and Motion Sickness in Mice. Front Neurol 9:918.

Isgrig K, Shteamer JW, Belyantseva IA, Drummond MC, Fitzgerald TS, Vijayakumar S, Jones SM, Griffith AJ, Friedman TB, Cunningham LL, Chien WW (2017) Gene Therapy Restores Balance and Auditory Functions in a Mouse Model of Usher Syndrome. Molecular Therapy 25:780–791.

Jamali M, Chacron MJ, Cullen KE (2016) Self-motion evokes precise spike timing in the primate vestibular system. Nat Commun 7:13229.

Jáuregui E, Scheinman K, BibriescaMejia I, Finkbeiner C, Phillips JA, Gantz JA, Phillips J, Stone J (2024) Sensorineural correlates of failed functional recovery after natural regeneration of vestibular hair cells in adult mice. Front Neurol.

Jones TA, Jones SM, Vijayakumar S, Brugeaud A, Bothwell M, Chabbert C (2011) The adequate stimulus for mammalian linear vestibular evoked potentials (VsEPs). Hear Res 280:133–140.

Kawamoto K, Izumikawa M, Beyer LA, Atkin GM, Raphael Y (2009) Spontaneous hair cell regeneration in the mouse utricle following gentamicin ototoxicity. Hear Res 247:17– 26.

Lahlou G, Calvet C, Simon F, Michel V, Alciato L, Plion B, Boutet De Monvel J, Lecomte M-J, Beraneck M, Petit C, Safieddine S (2024a) Extended time frame for restoring inner ear function through gene therapy in Usher1G preclinical model. JCI Insight 9:1–14.

Lahlou H, Zhu H, Zhou W, Edge ASB (2024b) Pharmacological regeneration of sensory hair cells restores afferent innervation and vestibular function. Journal of Clinical Investigation Available at: http://www.jci.org/articles/view/181201.

Lentz JJ, Jodelka FM, Hinrich AJ, McCaffrey KE, Farris HE, Spalitta MJ, Bazan NG, Duelli DM, Rigo F, Hastings ML (2013) Rescue of hearing and vestibular function by antisense oligonucleotides in a mouse model of human deafness. Nat Med 19:345–350.

Llorens J, Demêmes D, Sans A (1993) The behavioural syndrome caused by 3’3-Iminodipropionitrile and related nitriles in the rat is associated with degeneration of the vestibular sensory hair cells. Toxicology and Applied Pharmacology:199–210.

Lopez I, Ishiyama G, Tang Y, Tokita J, Baloh RW, Ishiyama A (2005) Regional estimates of hair cells and supporting cells in the human crista ampullaris. J Neurosci Res 82:421–431.

Lysakowski A, Goldberg JM (1997) A regional ultrastructural analysis of the cellular and synaptic architecture in the chinchilla cristae ampullares. J Comp Neurol 389:419– 443.

Mackowetzky K, Yoon KH, Mackowetzky EJ, Waskiewicz AJ (2021) Development and evolution of the vestibular apparatuses of the inner ear. J Anat 239:801–828.

Maroto AF, Barrallo-Gimeno A, Llorens J (2021) Relationship between vestibular hair cell loss and deficits in two anti-gravity reflexes in the rat. Hearing Research 410:108336.

Martins-Lopes V, Bellmunt A, Greguske EA, Maroto AF, Boadas-Vaello P, Llorens J (2019) Quantitative Assessment of Anti-Gravity Reflexes to EvaluateVestibular Dysfunction in Rats. JARO 20:553–563.

Maudoux A, Vitry S, El-Amraoui A (2022) Vestibular Deficits in Deafness: Clinical Presentation, Animal Modeling, and Treatment Solutions. Front Neurol 13:816534.

McInturff S, Burns JC, Kelley MW (2018) Characterization of spatial and temporal development of Type I and Type II hair cells in the mouse utricle using new cell-type-specific markers. Biology Open 7:bio038083.

Merchant SN, Tsuji K, Wall C (2000) Temporal Bone Studies of the Human Peripheral Vestibular System. Ann Otol Rhinol Laryngol.

Oommen BS, Stahl JS (2008) Eye orientation during static tilts and its relationship to spontaneous head pitch in the laboratory mouse. Brain Research 1193:57–66.

Paplou V, Schubert NMA, Pyott SJ (2021) Age-Related Changes in the Cochlea and Vestibule: Shared Patterns and Processes. Front Neurosci 15:680856.

Rauch SD, Velazquez-Villaseñor L, Dimitri PS, Merchant SN (2001) Decreasing Hair Cell Counts in Aging Humans. Annals of the New York Academy of Sciences 942:220–227.

Rüsch A, Lysakowski A, Eatock RA (1998) Postnatal development of type I and type II hair cells in the mouse utricle: acquisition of voltage-gated conductances and differentiated morphology. J Neurosci 18:7487–7501.

Sayyid ZN, Wang T, Chen L, Jones SM, Cheng AG (2019) Atoh1 Directs Regeneration and Functional Recovery of the Mature Mouse Vestibular System. Cell Rep 28:312–324.e4.

Schenberg L, Palou A, Simon F, Bonnard T, Barton C-E, Fricker D, Tagliabue M, Llorens J, Beraneck M (2023) Multisensory gaze stabilization in response to subchronic alteration of vestibular type I hair cells. eLife 12:RP88819.

Schlecker C, Praetorius M, Brough DE, Presler RG, Hsu C, Plinkert PK, Staecker H (2011) Selective atonal gene delivery improves balance function in a mouse model of vestibular disease. Gene Ther 18:884–890.

Schott JW, Huang P, Morgan M, Nelson-Brantley J, Koehler A, Renslo B, Büning H, Warnecke A, Schambach A, Staecker H (2023) Third-generation lentiviral gene therapy rescues function in a mouse model of Usher 1B. Molecular Therapy 31:3502–3519.

Selimoğlu E, Kalkandelen S, Erdoğan F (2003) Comparative vestibulotoxicity of different aminoglycosides in the Guinea pigs. Yonsei Med J 44:517–522.

Simon F, Pericat D, Djian C, Fricker D, Denoyelle F, Beraneck M (2020) Surgical techniques and functional evaluation for vestibular lesions in the mouse: unilateral labyrinthectomy (UL) and unilateral vestibular neurectomy (UVN). J Neurol 267:51– 61.

Simon F, Tissir F, Michel V, Lahlou G, Deans M, Beraneck M (2021) Implication of Vestibular Hair Cell Loss of Planar Polarity for the Canal and Otolith-Dependent Vestibulo-Ocular Reflexes in Celsr1–/– Mice. Front Neurosci 15:750596.

Soler-Martin C, Diez-Padrisa N, Boadas-Vaello P, Llorens J (2006) Behavioral Disturbances and Hair Cell Loss in the Inner Ear Following Nitrile Exposure in Mice, Guinea Pigs, and Frogs. Toxicological Sciences 96:123–132.

Songer JE, Eatock RA (2013) Tuning and Timing in Mammalian Type I Hair Cells and Calyceal Synapses. Journal of Neuroscience 33:3706–3724.

Stahl JS (2004) Using eye movements to assess brain function in mice. Vision Research 44:3401–3410.

Walther LE, Westhofen M (2008) Presbyvertigo-aging of otoconia and vestibular sensory cells. VES 17:89–92.

Wang T, Chai R, Kim GS, Pham N, Jansson L, Nguyen D-H, Kuo B, May LA, Zuo J, Cunningham LL, Cheng AG (2015) Lgr5+ cells regenerate hair cells via proliferation and direct transdifferentiation in damaged neonatal mouse utricle. Nat Commun 6:6613.

Wang T, Niwa M, Sayyid ZN, Hosseini DK, Pham N, Jones SM, Ricci AJ, Cheng AG (2019) Uncoordinated maturation of developing and regenerating postnatal mammalian vestibular hair cells. PLoS Biol 17:e3000326.

Wu P, Wu X, Zhang C, Chen X, Huang Y, Li H (2021) Hair Cell Protection from Ototoxic Drugs. Neural Plast 2021:4909237.

Zakir M, Dickman JD (2006) Regeneration of Vestibular Otolith Afferents after Ototoxic Damage. Journal of Neuroscience 26:2881–2893.

Zeng S, Ni W, Jiang H, You D, Wang J, Lu X, Liu L, Yu H, Wu J, Chen F, Li H, Wang Y, Chen Y, Li W (2020) Toxic Effects of 3,3 1 -Iminodipropionitrile on Vestibular System in Adult C57BL/6J Mice In Vivo. Neural Plasticity 2020:1–11.

