## Supplementary figures and images for "Hair cell population integrity necessary to preserve vestibular function"

### Supplementary Figure 1

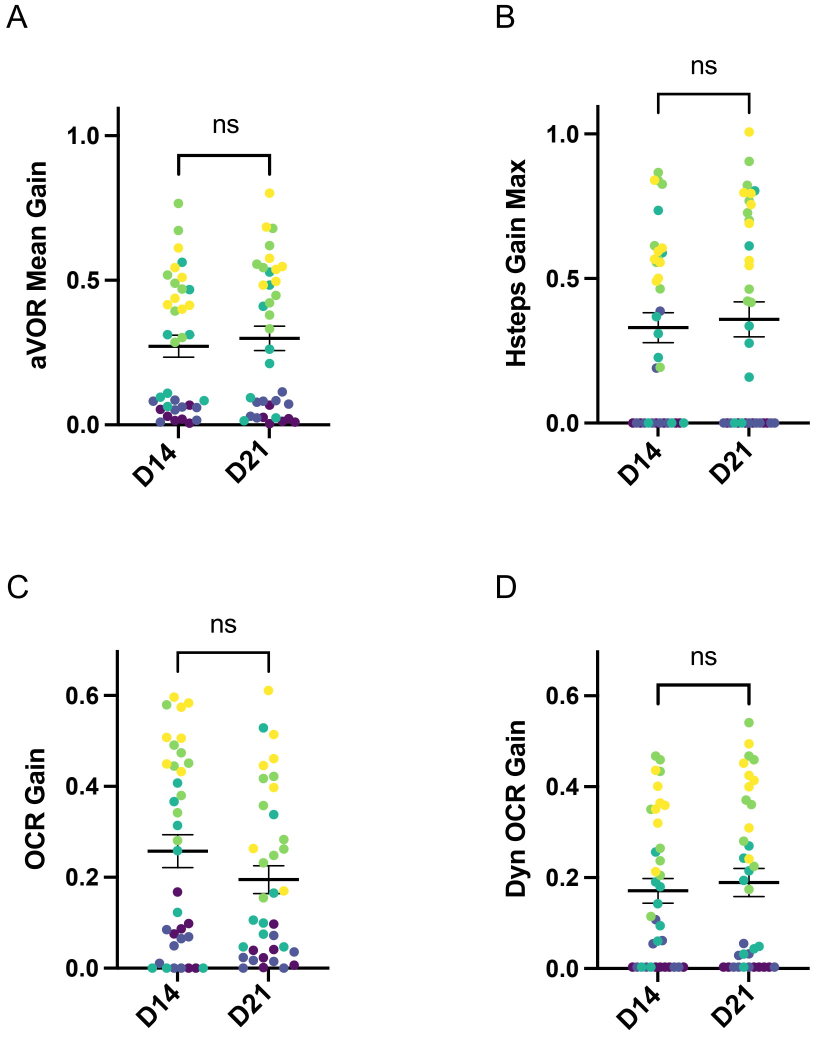

### Supplementary Figure 2

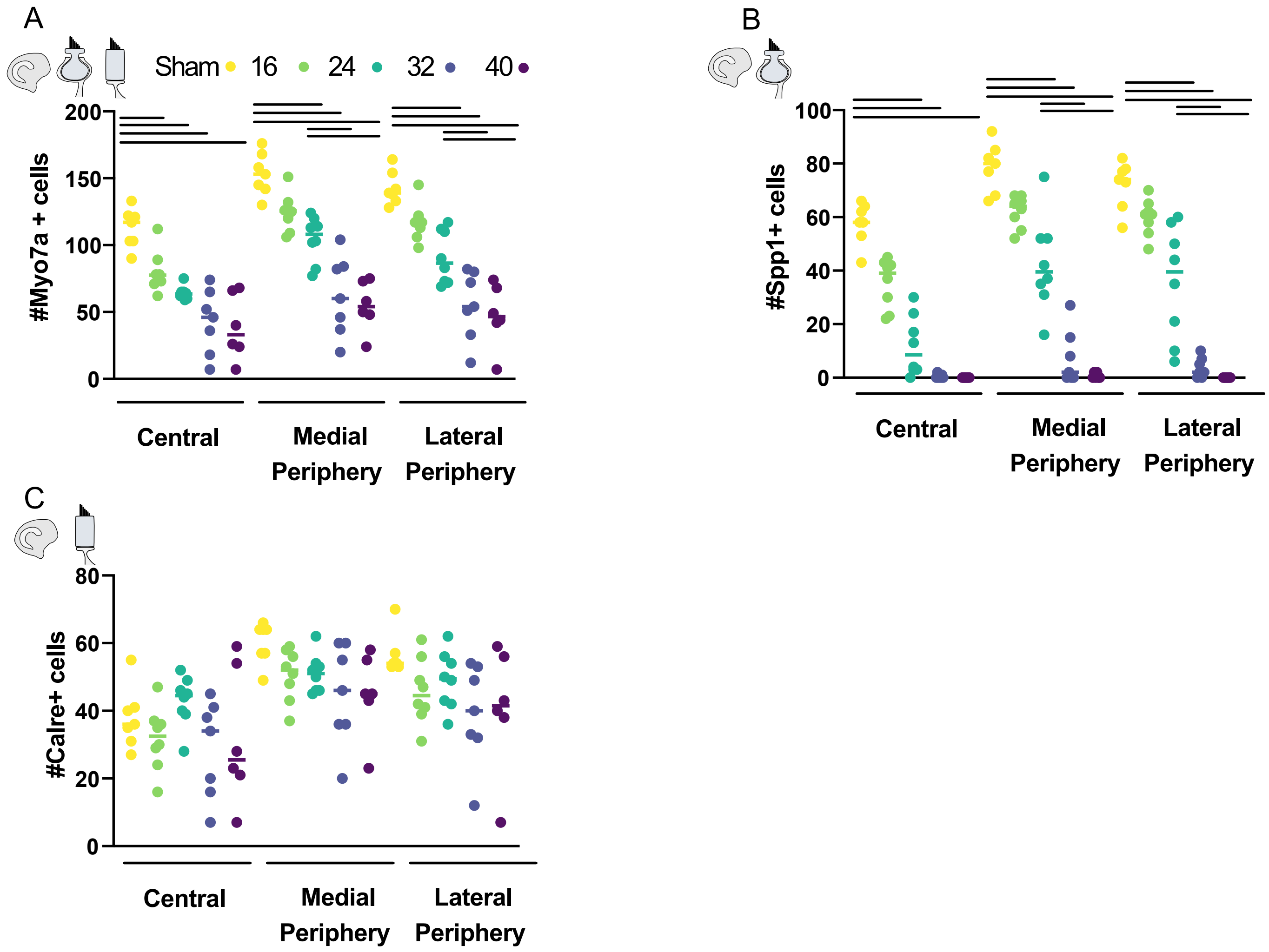
